# Acquisition and repeated alteration of (TTGGG)_n_ telomeric repeats in Odonata (dragonflies and damselflies)

**DOI:** 10.1101/2025.05.20.653629

**Authors:** Tatsuhiro Gotoh, Haruka Suzuki, Minoru Moriyama, Ryo Futahashi, Mizuko Osanai-Futahashi

## Abstract

Chromosome ends of most eukaryotes are composed of simple telomeric repeats. For arthropods, TTAGG pentanucleotide repeats, (TTAGG)_n_ has been considered as the ancestral telomeric repeat. However, in the order Odonata, the earliest diverged group in insects that contains dragonflies and damselflies, (TTAGG)_n_ signal has been almost undetectable in most examined species. Here, we report a unique pentanucleotide repeat (TTGGG)_n_ as the typical telomeric repeat sequence for Odonata (dragonflies and damselflies). Based on genomic information from ten Odonata species, (TTGGG)_n_ was considered the most likely candidate for telomeric repeat sequences. By fluorescence *in situ* hybridization (FISH) using 12 Odonata species, clear (TTGGG)_n_ signals were detected at the chromosome ends in both dragonflies and damselflies. By Southern hybridization using 63 Odonata species, strong (TTGGG)_n_ signals were detected from the majority of species covering all three suborders of Odonata, indicating that the telomeric repeat of the common ancestor of extant Odonata is (TTGGG)n. Notably, there were a few Odonata species in which (TTGGG)n signals were faint or absent, suggesting that the telomeric repeat sequence has repeatedly diverged in Odonata, even within genera such as *Sympetrum*. We also identified telomerase genes in both dragonflies and damselflies, and in some species, more than two telomerase genes are suggested to be present. Overall, this study demonstrates the ancestral acquisition of novel telomeric repeats and their repeated alteration in Odonata.

## 1. Introduction

The ends of linear eukaryotic chromosomes, termed telomeres, are generally composed of simple repeated sequences. One strand of the repeats is G-rich with its 3′ -end pointing to the chromosome end. The G-rich strand is synthesized by the enzyme telomerase, which is composed of telomerase reverse transcriptase (TERT) and template RNA (Blackburn, 1991). Since chromosome ends cannot be replicated due to the end replication problem, the addition of telomeric repeats by telomerase is crucial for many organisms to prevent shortening of chromosome ends.

The common pattern of telomeric repeat sequences across eukaryotes is (TxAyGz)_n_. For example, (TTAGGG)_n_ and (TTTAGGG)_n_ are the telomeric repeat widespread in metazoans and plants, respectively (Traut et al., 2007; Peska and Garcia, 2020). (TTAGG)_n_, first discovered as the telomeric repeat sequence of the silkworm *Bombyx mori* (Okazaki et al., 1993), has been considered as the ancestral telomeric repeat sequence for arthropods (Okazaki et al., 1993; Sahara et al., 1999; Frydrychová et al., 2004; Vítková et al., 2005; Traut et al., 2007). However, (TTAGG)_n_ cannot be detected in several insect orders, i.e. Diptera, Mecoptera, Dermaptera, Ephemeroptera, and Odonata (Frydrychová et al., 2004; Mason et al., 2016; Kuznetsova et al., 2021). In Diptera, telomerase reverse transcriptase (TERT) gene is absent in the genome, and the chromosome ends are maintained by alternative systems such as telomere specific retrotransposons HeT-A and TART in *Drosophila melanogaster*, and recombination in mosquitos (Roth et al., 1997; Mason et al., 2016).

Alteration of telomeric repeat sequence has been reported within several insect orders. In Coleoptera, the replacement of (TTAGG)_n_ by (TCAGG)_n_ has been detected in superfamilies Tenebrionoidea and Cleroidea, and within Curculionoidea and Chrysomeloidea, and also by (TTGGG)n in *Anoplotrupes stercorosus* (Geotrupidae, Scarabaeoidea) (Osanai et al., 2006; Tribolium Genome Sequencing Consortium, 2008; Osanai-Futahashi and Fujiwara, 2011; Mravinac et al., 2011; Prušáková et al., 2021). In Hymenoptera, (TTAGG)_n_ telomeric repeats have been repeatedly replaced, and a variety of telomeric repeat sequences have been reported (e.g. (TTATTGGG)_n_, (TTAGGTTGGGG)_n_, (TTAGGTCTGGG)_n_, (TTAGGG)_n_, (TTGCGTCTGGG)_n_, (TTAGGTCTGGG)_n_) (Zhou et al., 2022; Fajkus et al., 2023; Lukhtanov and Pazhenkova, 2023). In Hemiptera, (TTAGG)_n_ is lost in many lineages, and (TTTTAGGGATGG)n, (TTAGGGGTGG)n, (TTAGGGGTGGT)_n_ are detected in Pentatomomorpha (Sahara et al., 1999; Lukhtanov and Pazhenkova, 2023; Stoianova et al., 2024).

Absence of (TTAGG)_n_ has been also reported in Odonata species (Table S1) (Frydrychova et al., 2004; Kuznetsova et al., 2017; Kuznetsova et al., 2021). In one dragonfly species *Sympetrum vulgatum*, (TTAGG)_n_ has been detected by fluorescence *in situ* hybridization (FISH), although the (TTAGG)_n_ signals were detected in the center of the cruciform structured meiotic chromosomes, suggesting that signal locations are not at the chromosome ends (Kuznetsova et al., 2021). In one damselfly species *Platycnemis pennipes*, (TTGGG)_n_ repeat sequence has been reported in genome sequence (Lukhtanov and Pazhenkova, 2023), however, the chromosomal location of this sequence has not been confirmed by FISH. Here, we report detection of (TTGGG)_n_ repeat sequence in the chromosome ends of various Odonata species by FISH as well as genome sequence analysis. The widespread distribution of (TTGGG)_n_ in dragonflies and damselflies indicates that the (TTGGG)_n_ is the typical and ancestral telomeric repeat sequence of Odonata. Meanwhile, Southern hybridization and genome sequence analysis indicates that the alteration of (TTGGG)_n_ have occurred repeatedly.

## 2. Materials and methods

### 2.1. Repeat extraction by repeat masker

Repeats were extracted using RepeatModeler 2.0.5 and RepeatMasker 4.1.6 software (https://repeatmasker.org/) (Flynn et al., 2020) from genome sequences of the following ten Odonata species registered in the National Center for Biotechnology Information (NCBI) database; *Tanypteryx hageni* (accession no. GCA_028673005.1), *Pantala flavescens* (GCA_020796165.1)*, Sympetrum striolatum* (GCA_947579665.1)*, Sympetrum sanguineum* (GCA_964198035.1), *Pyrrhosoma nymphula* (GCA_963573305.1)*, Ceriagrion tenellum* (GCA_963169105.1), *Ischnura elegans* (GCA_921293095.1)*, Ischnura senegalensis* (GCA_964251815.1), *Platycnemis pennipes* (GCA_933228895.1), and *Hetaerina titia* (GCA_037158775.1). The repeats classified as simple repeats were examined for sequence divergence, length and their chromosomal location.

### 2.2. Chromosome preparations and fluorescence *in situ* hybridization

Alexa488-OO(CCCAA)3CCC and Alexa488-OO(CCTAA)3CCT Peptide nucleic acid (PNA) probes were synthesized by PANAGENE. Odonata specimens used for FISH were collected in Ibaraki prefecture. Chromosome preparations were prepared from gonads or embryos. To collect dragonfly eggs, the females were let to lay eggs in glass vials filled with water. Eggs were transferred onto a slide, briefly air dried, and the dried jelly coat was removed by forceps. Chromosome preparation of Odonata species other than *Pantala flavescens* and *Orthetrum albistylum* were performed as follows. Embryos or gonads were dissected in Phosphate buffered saline (PBS), incubated in a hypotonic solution (75 mM KCl) for 10 minutes, and then fixed in 100% ethanol : glacial acetic acid = 3 : 1 at 4□ for one day or more. The fixed samples were placed on a slide, and covered with a drop of 45% acetic acid, then physically squashed using a coverslip. Subsequently, the slide/coverslip was flash frozen in liquid nitrogen. After carefully removing slides from liquid nitrogen, coverslips were removed with a razor blade. After fixation, slides were washed thrice in 0.5% Tween-20 in PBS and dehydrated in subjected to an ethanol series (70%, 90%, 100%, 3 minutes each), and rinsed with hybridization buffer (30% formamide in PBS). Slides were then denatured at 92□ for 5 minutes, and hybridization was performed at 37□ for overnight with hybridization solution containing 0.17 µM PNA probe and 3 % salmon sperm DNA. The slides were washed twice with hybridization wash buffer (30% formamide in 10 mM Tris-HCl) for 15 minutes, once each by 2 x SSC and 0.1 x SSC for 5 minutes. Then the slides were subjected to an ethanol series (70%, 90%, 100%, 3 min each), stained in 2 µg/ml DAPI (Dojindo) or Propidium iodide (Dojindo) in PBS for 30 minutes, washed thrice with PBS for 5 minutes, and mounted in Vectashield mounting medium (Vector laboratories). Fluorescence signals were observed under Zeiss Axio Observer inverted microscope, and images were taken with Axiocam 503 mono camera (Carl Zeiss). Zen software (Carl Zeiss) was used to control the system. Images of *R. fuliginosa*, *S. frequens*, *S. baccha* were taken under same exposure conditions and the signals of Alexa488-OO(CCCAA)3CCC were analyzed by ImageJ.

For *Pantala flavescens*, dissected embryos were placed on a new slide, and excess PBS was removed. The embryos and gonads were then physically squashed using a coverslip. Subsequently, the slide/coverslip were flash frozen in liquid nitrogen. After carefully removing slides from liquid nitrogen, coverslips were removed with a razor blade. Embryos were fixed for 3 minutes in freshly prepared Carnoy fixative (ethanol:chloroform:acetic acid, 6:3:1). The procedure after fixation is the same as above.

For *Orthetrum albistylum*, eggs were shaked in Heptane : 5% HCHO in PHEM buffer (4 mM MgSO_4_, 30 mM HEPES, 60 mM PIPES, 10 mM EGTA) = 1 : 1 at 500 rpm for 3 minutes using Mix-VR2 (Taitec). The embryos were dissected out of the eggs in PBS with forceps and fine insect pins and placed on a slide, and post-fixed in methanol for 5 minutes. After fixation, slides were washed thrice in 0.5% TritonX-100 in PBS and subjected to an ethanol series (70%, 90%, 100%, 3 minutes each), and rinsed with hybridization buffer (30% formamide in PBS). Slides were then denatured at 85□ for 5 minutes, and hybridization was performed at 37□ for 2 hours in hybridization solution containing 0.17 µM PNA probe and 3 % salmon sperm DNA. The procedure after hybridization is the same as other Odonata species.

To prepare *Bombyx mori* chromosome preparation, ovaries were dissected from day-2 larva of the 5th instar *w1, pnd* strain, and incubated in a hypotonic solution (75 mM KCl) for 10 minutes. After fixation in 1% paraformaldehyde in PHEM buffer for 6 minutes, the ovary was placed on a MAS coated slide (MATSUNAMI), and then physically squashed using a coverslip. Subsequently, the slide/coverslip was flash frozen in liquid nitrogen. After carefully removing slides from liquid nitrogen, coverslips were removed with a razor blade. The ovaries were post-fixed with 100% methanol : glacial acetic acid = 3 : 1 for 10 minutes. After fixation, slides were washed thrice in 0.5% TritonX-100 in PBS and subjected to an ethanol series (70%, 90%, 100%, 3 minutes each), and rinsed with hybridization buffer (30% formamide in PBS). Slides were then denatured at 85□ for 5 minutes, and hybridization was performed at 37□ for 2 hours in hybridization solution containing 0.17 µM PNA probe and 3 % salmon sperm DNA. Rest of the procedure is the same as FISH of Odonata specimen.

### 2.3. Southern hybridization

Hybridization probes were prepared using non-template PCR, using TaKaRa ExTaq (TaKaRa). The reaction included digoxigenin 11-dUTP (Roche) to label the PCR products. The primers used for generating TTAGG probe are (TTAGG)_6_ and (CCTAA)_6_, and primers used for TTGGG probe are (TTGGG)_6_ and (CCCAA)_6_. Specimen used for genome extraction is described in Table S2. Genomic DNAs were extracted using Maxwell 16 Tissue DNA Purification kit (Promega), or PureLink ^TM^ Genomic DNA Mini kit (Invitrogen). From each species, about 1 μg genomic DNA was digested with HindIII (TaKaRa), separated on 0.8% agarose gel, and blotted onto a Hybond-N+ nylon membrane (Amersham Biosciences) by capillary transfer in 20× SSC overnight, followed by Southern with chemiluminescence detection as described previously (Traut et al., 2007), with slight modifications. Briefly, prehybridization was conducted with 5 ml of DIG Easy Hyb (Roche) for 30min, and the hybridization was performed with DIG Easy Hyb containing DIG-labeled probe (25 ng/ml), at 52 °C for 3 h and 42 °C for 16 h for TTGGG probe and TTAGGG probe, respectively. Washes and blocking were performed using DIG Wash and Block Buffer Set (Roche), according to the manufacturer’s protocol. The chemiluminescence detection was performed using the alkaline phosphatase-CDP-Star system (Roche) and the signals were detected using the CCD camera LAS-3000 (Fujifilm).

### 2.4. Transcriptome analyses

Transcriptome analyses were performed as described (Futahashi et al., 2019). Total RNA was extracted from the freshly prepared samples using Maxwell 16 LEV Simply RNA Tissue kit (Promega), and cDNA libraries were constructed using TruSeq RNA Sample Preparation Kits v2 (Illumina) or NEBNext Ultra II Directional RNA Library Prep Kit for Illumina (NEB) and sequenced by HiSeq (Illumina). The accession numbers of RNA-sequencing data used in this study are listed in Table S3.

The raw reads were subjected to de novo assembly using the Trinity program v. 2.4.0 (Grabherr et al. 2011). After automatic assembling, we checked and manually corrected the sequences of telomerase gene by using the Integrative Genomics Viewer (Thorvaldsdóttir et al. 2013).

### 2.5. Phylogenetic analysis

Deduced amino acid sequences of TERT genes were aligned using the Clustal W program implemented in the program package MEGA 11 (Tamura et al., 2021). Molecular phylogenetic analyses were conducted by neighbor-joining method and the maximum-likelihood method using MEGA11. Bootstrap values were obtained by 1,000 resampling.

## 3. Results

### 3.1. Detection of (TTGGG)n as a candidate for telomeric repeats in genomic sequence of odonata

To find candidate telomeric repeat sequences of Odonata, we extracted repeats using RepeatMasker software (Flynn et al., 2020) from the genome sequences of ten Odonata species (one Petaluridae species, three Libellulidae species, four Coenagrionidae species, one Platycnemididae species, and one Calopterygidae species) whose genomes were assembled at the chromosome level (Table 1, Supplementary data 1-10). Petaluridae and Libellulidae belong to the suborder Anisoptera, while Coenagrionidae, Platycnemididae, and Calopterygidae belong to the suborder Zygoptera.

**Table 1.**
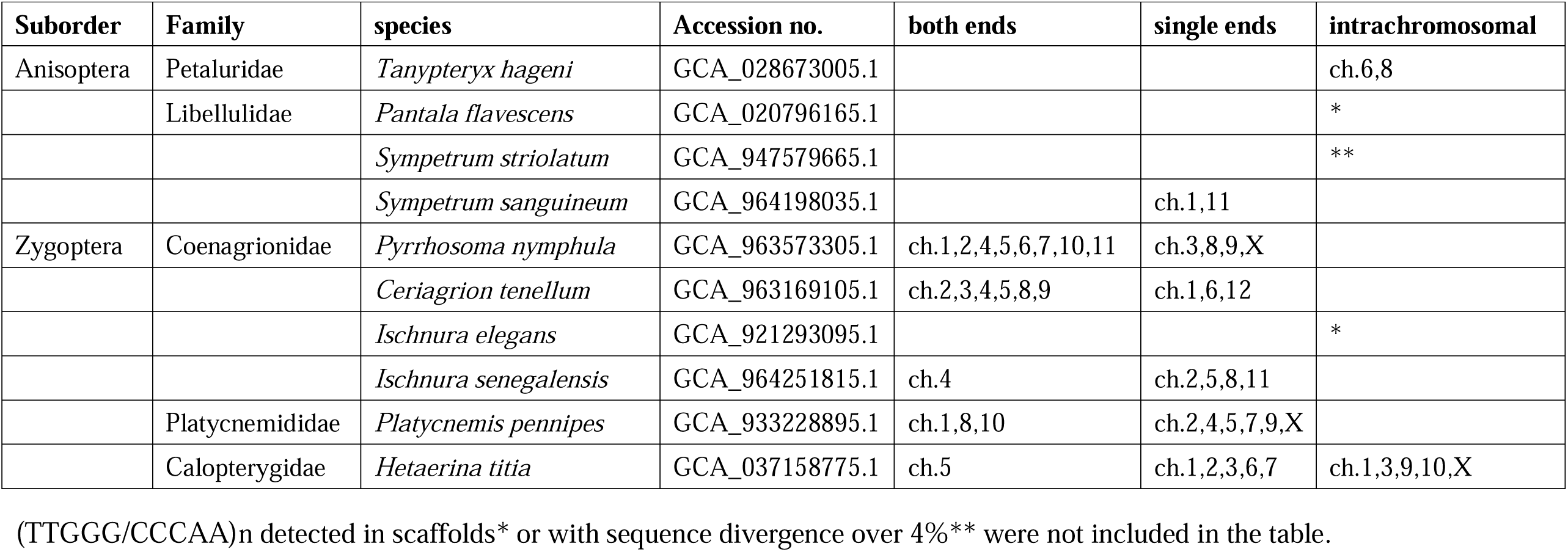
Survey for (TTGGG/CCCAA)n repeats in the publicly available Odonata genomes.

We focused on repeats classified into “simple repeats”, and searched for long repeats with low sequence divergence that were located on the ends of the chromosome assembly. In three damselflies *P. nymphula*, *C. tenellum*, and *P. pennipes*, pentanucleotide repeats (TTGGG)n and its reverse complement (CCCAA)n of low sequence divergence (mostly under 1%) ranging from 1 kb to 8.6 kb were found in ends of the majority of chromosomes (Table 1; Supplementary data 5, 6, 9), suggesting that (TTGGG)n was the telomeric repeats of these species. For example, (TTGGG/CCCAA)n was found in both ends of the chromosomes 1, 2, 4, 5, 6, 7, 10, 11 and single ends of chromosomes 3, 8, 9, X in *P. nymphula* (Table 1; Supplementary data 5). (TTGGG/CCCAA)n was also found in chromosomal ends in *S. sanguineum* and *I. senegalensis* (Table 1; Supplementary data 4, 8). In *H. titia*, (TTGGG/CCCAA)n was found not only in chromosomal ends but also within intrachromosomal locations of chromosomes (Table 1; Supplementary data 10). In *P. flavescens* and *I. elegans*, (TTGGG/CCCAA)n with sequence divergence under 1% were found, but were shorter than 1.3 kb and were not located at the assembled chromosome termini (Table 1; Supplementary data 2, 7). Notably, (TTGGG/CCCAA)n of *Sympetrum striolatum* had higher sequence divergence (4 to 13%) than *Sympetrum sanguineum* (0 to 1.6%), and was not located at the chromosome ends (Supplementary data 3, 4). Exceptionally, one end of chromosome 11 of *C. tenellum* was 3.8 kb (TGGGC/GCCCA)n repeats (Supplementary data 6). Meanwhile, no stretch of other repeat sequences was found in the chromosome termini.

### 3.2. Detection of TTGGG repeats by FISH in the chromosome ends of Odonata species

Our survey with the NCBI genome sequence data indicated that (TTGGG)n is a strong candidate for the telomeric repeat sequence at least in some Odonata species. To investigate this possibility, we conducted fluorescence *in situ* hybridization (FISH) analyses. In vertebrates, PNA probe efficiently detects TTAGGG telomeric repeats in FISH (Genet et al., 2013). First, we performed FISH using (CCTAA)3CCT probe against the silkworm *B. mori*, whose telomeric repeat is known to be (TTAGG)_n._ As expected, clear (TTAGG)_n_ signals were detected at the chromosome ends (Fig. 1A). Next, we performed FISH using (CCCAA)3CCC probe against the Libellulidae dragonfly *O. albistylum*. Similarly, clear signals were visualized at the chromosome ends (Fig. 1B). Southern hybridization confirmed that the TTAGG and TTGGG probes were exclusively detected in *B. mori* and *O. albistylum*, respectively (Fig. 1C, 1D). These results demonstrate that (TTGGG)_n_ is a telomeric repeat sequence, at least in *O. albistylum*.

**Fig. 1.**
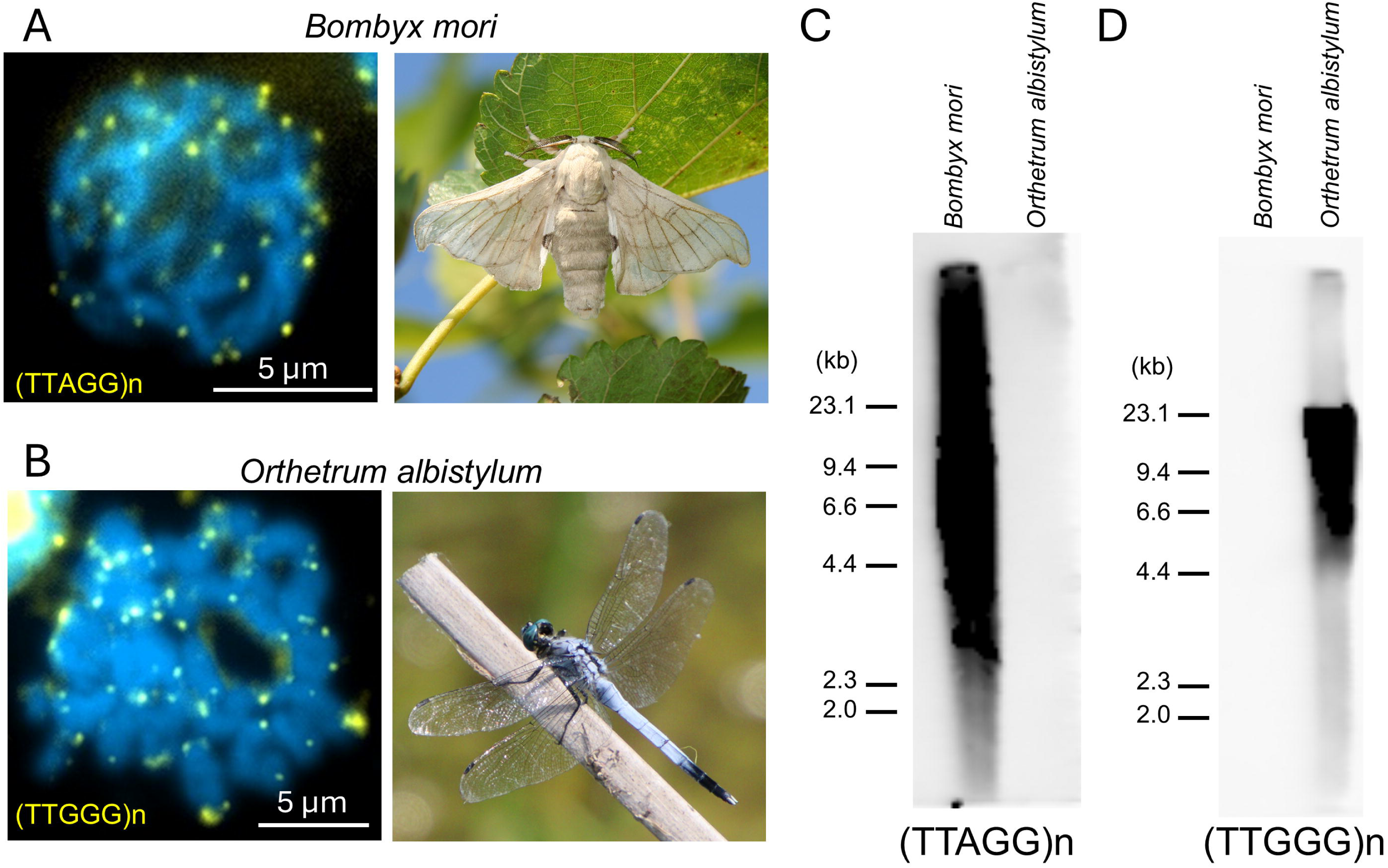
Detection of (TTAGG)n and (TTGGG)n by fluorescence *in situ* hybridization and southern hybridization and in the silkworm *Bombyx mori* and the dragonfly *Orthetrum albistylum.* (A) Fluorescence *in situ* hybridization of *B. mori* chromosomes using ovaries from 3^rd^ day of 5^th^ instar larvae, with (CCTAA)3CCT PNA probes. (B) Fluorescence *in situ* hybridization of *O. albistylum* chromosomes using embryo, with Alexa488-(CCCAA)3CCC PNA probes. Signals of probes are indicated in yellow, and chromosomes counter-stained with DAPI or PI are in blue. (C, D) Southern hybridization of *Bombyx mori* and *Orthetrum albistylum* genomes with probe targeting (TTAGG)n or (TTGGG)n.

To determine whether (TTGGG)_n_ is also a telomeric repeat sequence in other Odonata species, FISH was performed on additional eleven Odonata species including seven dragonfly species; *Anax parthenope* (Aeshnidae), *Anotogaster sieboldii* (Cordulegastridae), *Rhyothemis fuliginosa*, *Sympetrum frequens*, *Sympetrum baccha*, *Crocothemis servilia*, *Pantala flavescens* (Libellulidae), and four damselfly species; *Lestes temporalis* (Lestidae), *Aciagrion migratum*, *Ischnura senegalensis*, and *Ischnura asiatica* (Coenagrionidae). Nine of the eleven species, except for the two *Sympetrum* species, exhibited clear signals at the chromosomal ends (Figs. 2, S1, S2), as in *O. albistylum*. In chromosomes at the diakinesis stage, signals were detected at the telomeres at the tips of the chromosomal crosses (Fig. S1). In some cells, signals implying telomere bouquet structures with chromosomes looping in the same direction were also detected (Figs. 2H, S2). On the other hand, (TTGGG)_n_ signal intensity was weak in two *Sympetrum* species, especially in *S. frequens*, with only a faint signal detected (Figs. 2D, 2E, S3, Table S4).

**Fig. 2.**
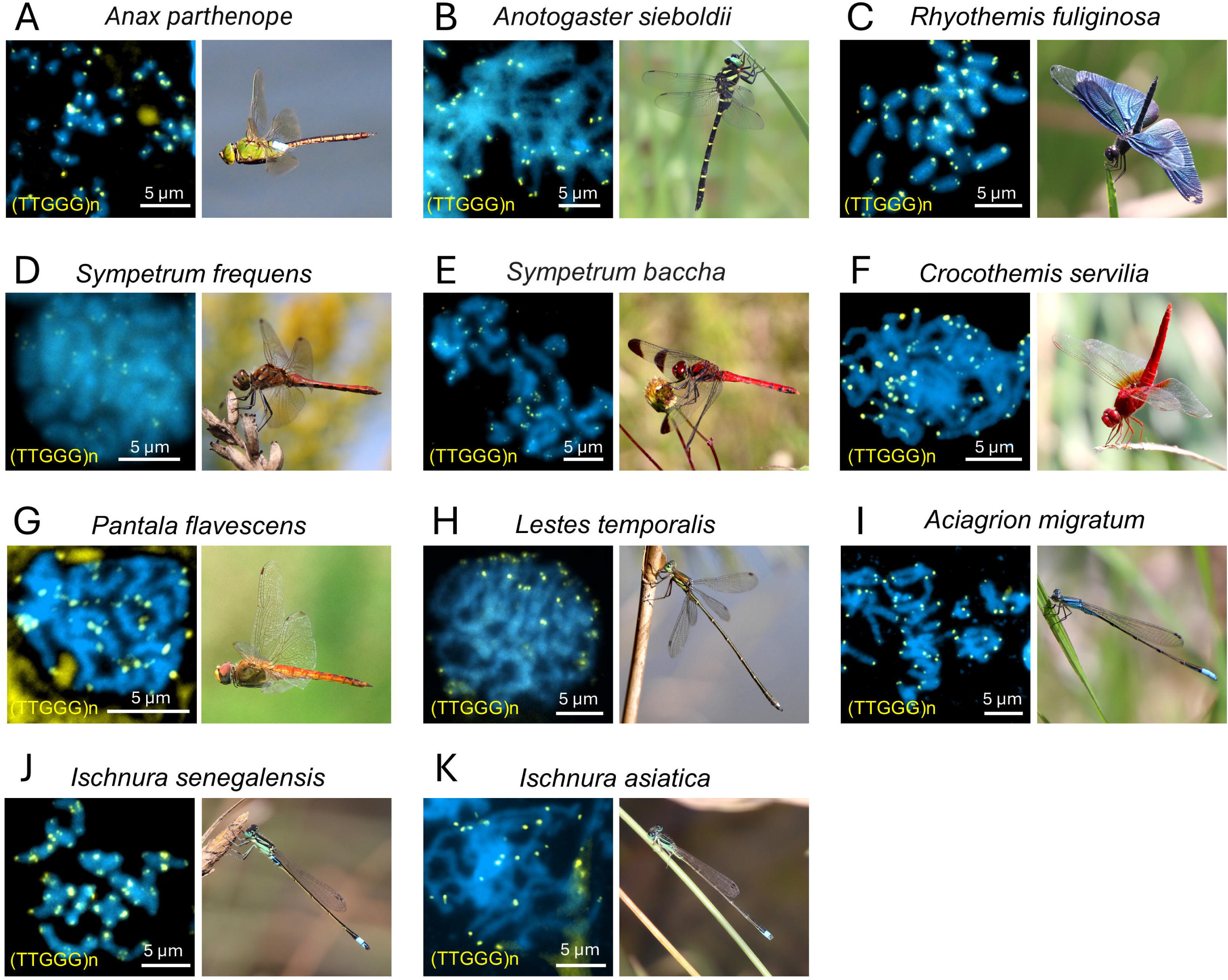
Fluorescence *in situ* hybridization of odonata species with a probe against (TTGGG)n. Alexa488-(CCCAA)3CCC PNA were used as probes. Signals of probes are indicated in yellow, and chromosomes counter-stained with DAPI or PI are in blue.

**Figure 3.**
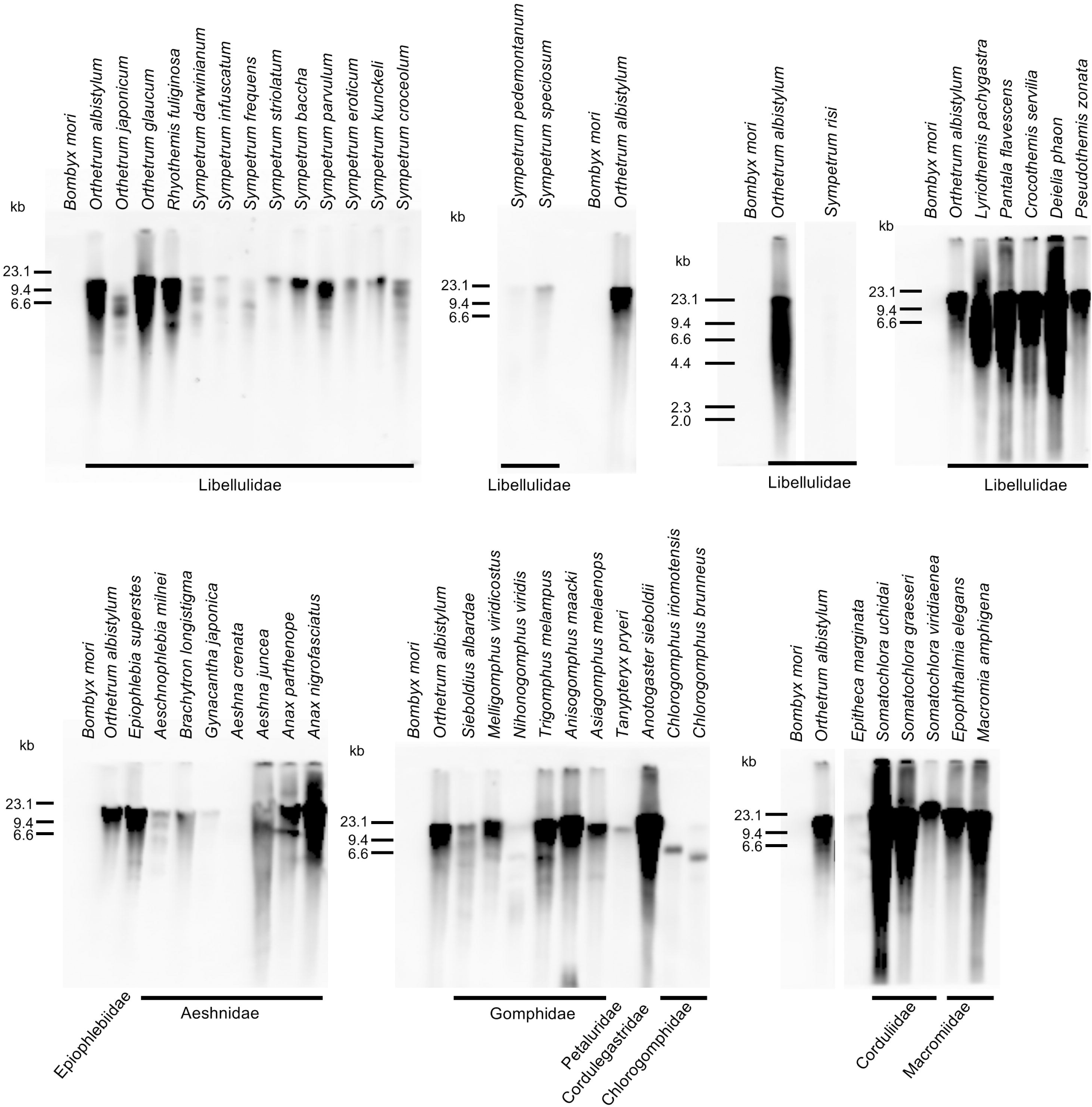
Screening of (TTGGG)n telomeric repeats of dragonfly species by Southern hybridization. The DIG-labeled probe specific to (TTGGG)n sequence was hybridized with HindIII-digested genomic DNAs of selected dragonfly (Anisoptera and Anisozygoptera) species. *Bombyx mori* genomic DNA was used as a negative control.

### 3.3 Distribution of TTGGG telomeric repeats across Odonata

The order Odonata consists of three suborders, Anisoptera (dragonflies), Zygoptera (damselflies), and Anisozygoptera (ancient dragonflies). To survey the distribution of (TTGGG)n telomeric repeats across Odonata species comprehensively, we performed Southern hybridization for 44 Anisoptera species (8 families), 18 Zygoptera species (5 families), and one Anisozygoptera species (Figs. 3, 4). *O. albistylum* and *B. mori* were used for positive and negative controls, respectively.

As a result, strong (TTGGG)n signals were detected from the majority of Odonata species, which covered all three suborders (Figs. 3, 4, 5, 6), indicating that the telomeric repeat of the common ancestor of extant Odonata is (TTGGG)n. It should be noted there were a few Odonata species, especially in the genus *Sympetrum*, which (TTGGG)n signals were faint or absent compared to *O. albistylum* (Figs. 3, 4, 5, 6). The amount of genome DNA used for these species excluding *Pseudocopera annulata*, were comparable to *O. albistylum* (Fig. S4), suggesting the alteration of their telomeric repeat sequences.

**Figure 4.**
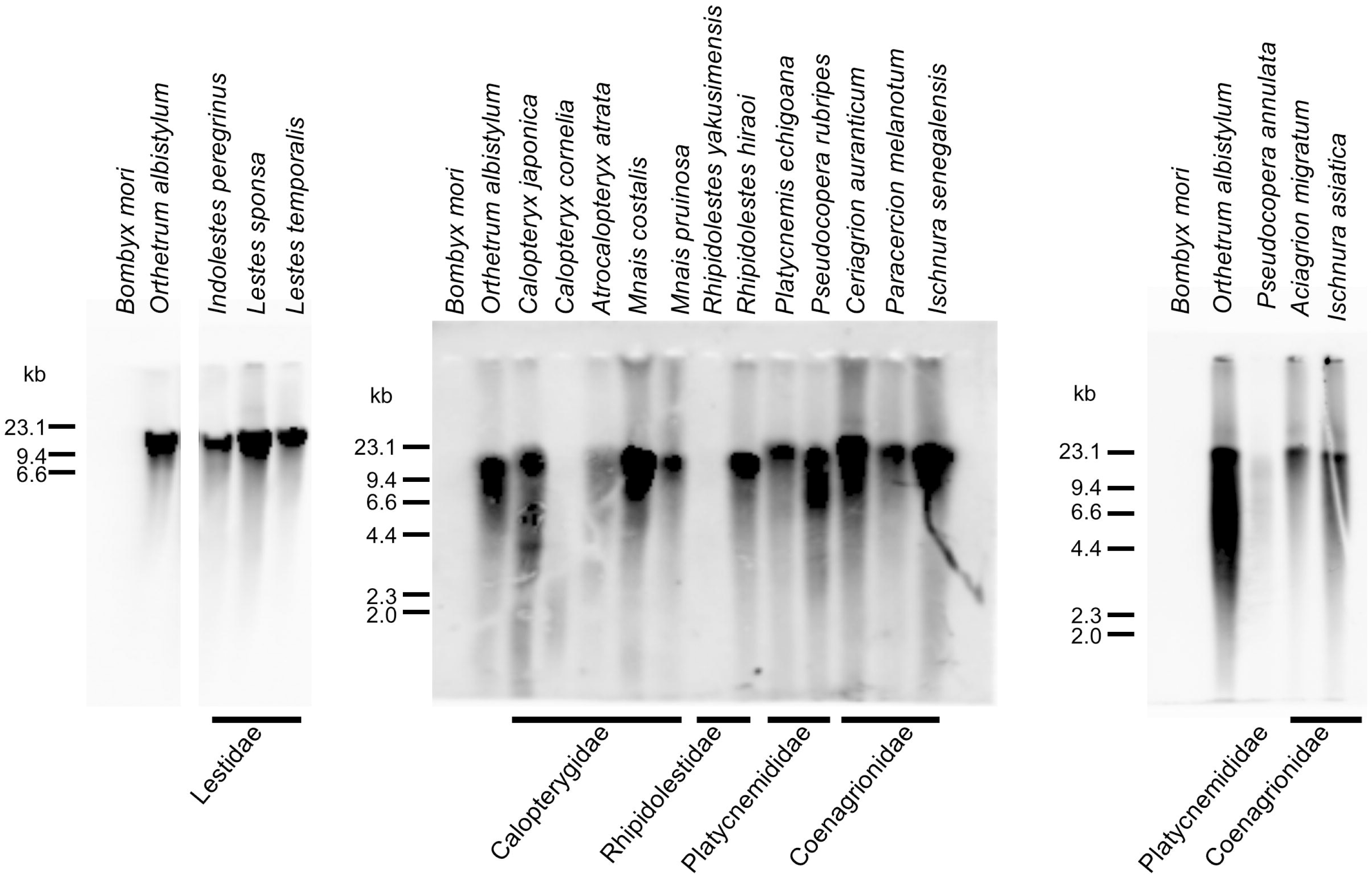
Screening of (TTGGG)n telomeric repeats of damselfly species by Southern hybridization. The DIG-labeled probe specific to (TTGGG)n sequence was hybridized with HindIII-digested genomic DNAs of selected damselfly (Zygoptera) species. *Bombyx mori* genomic DNA was used as a negative control.

**Figure 5.**
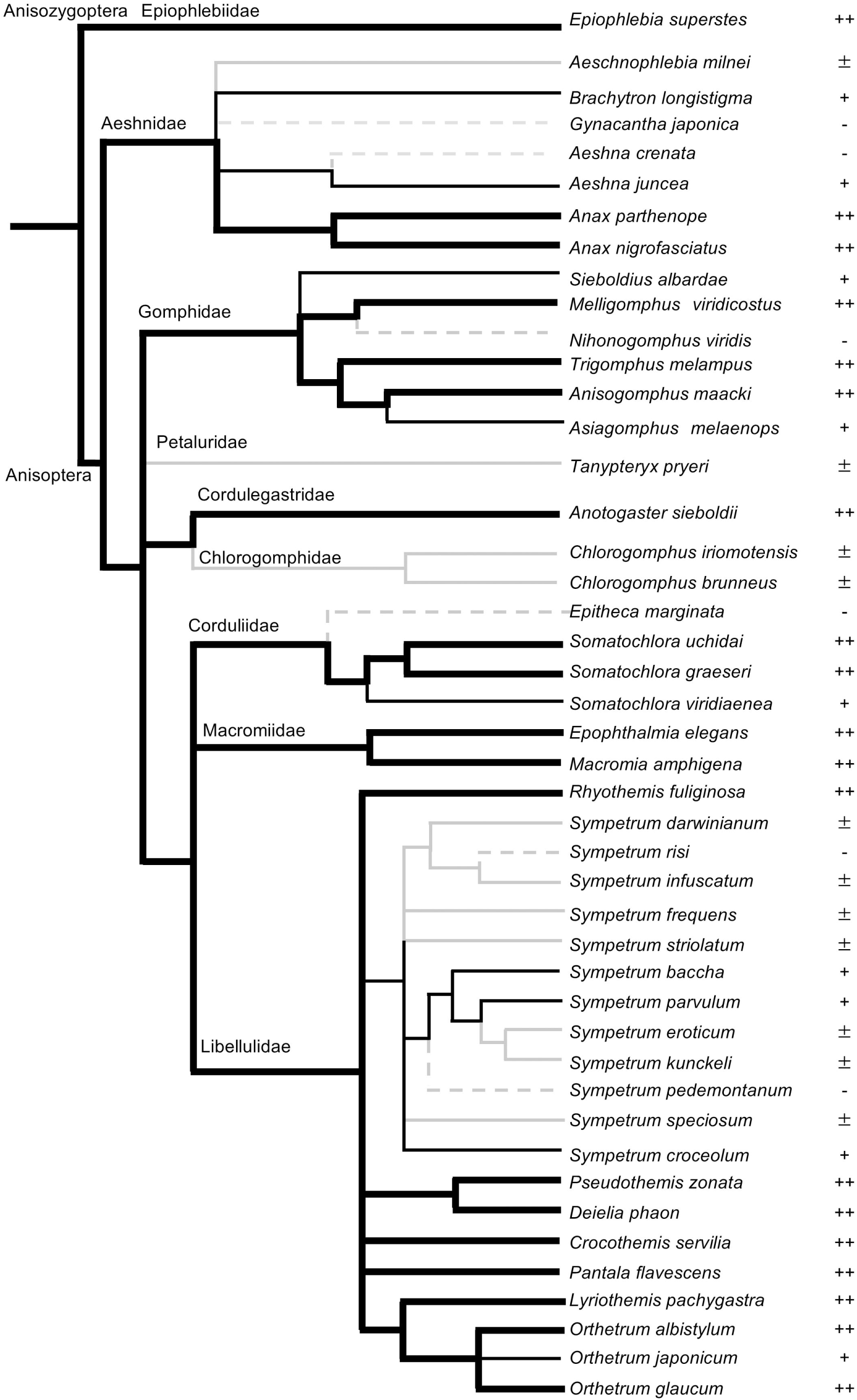
The presence of (TTGGG)_n_ telomeric sequence in dragonflies. The odonate phylogeny is based on Ozono et al., 2022. Southern hybridization signal intensity nearly equal to or greater than that of *Orthetrum albistylum* (++, thick black branch), weaker than that of *Orthetrum albistylum* but clearly visible (+, thin black branch); weak signal (+-, gray solid branch); nearly or no detectable signal (gray dotted branch).

**Figure 6.**
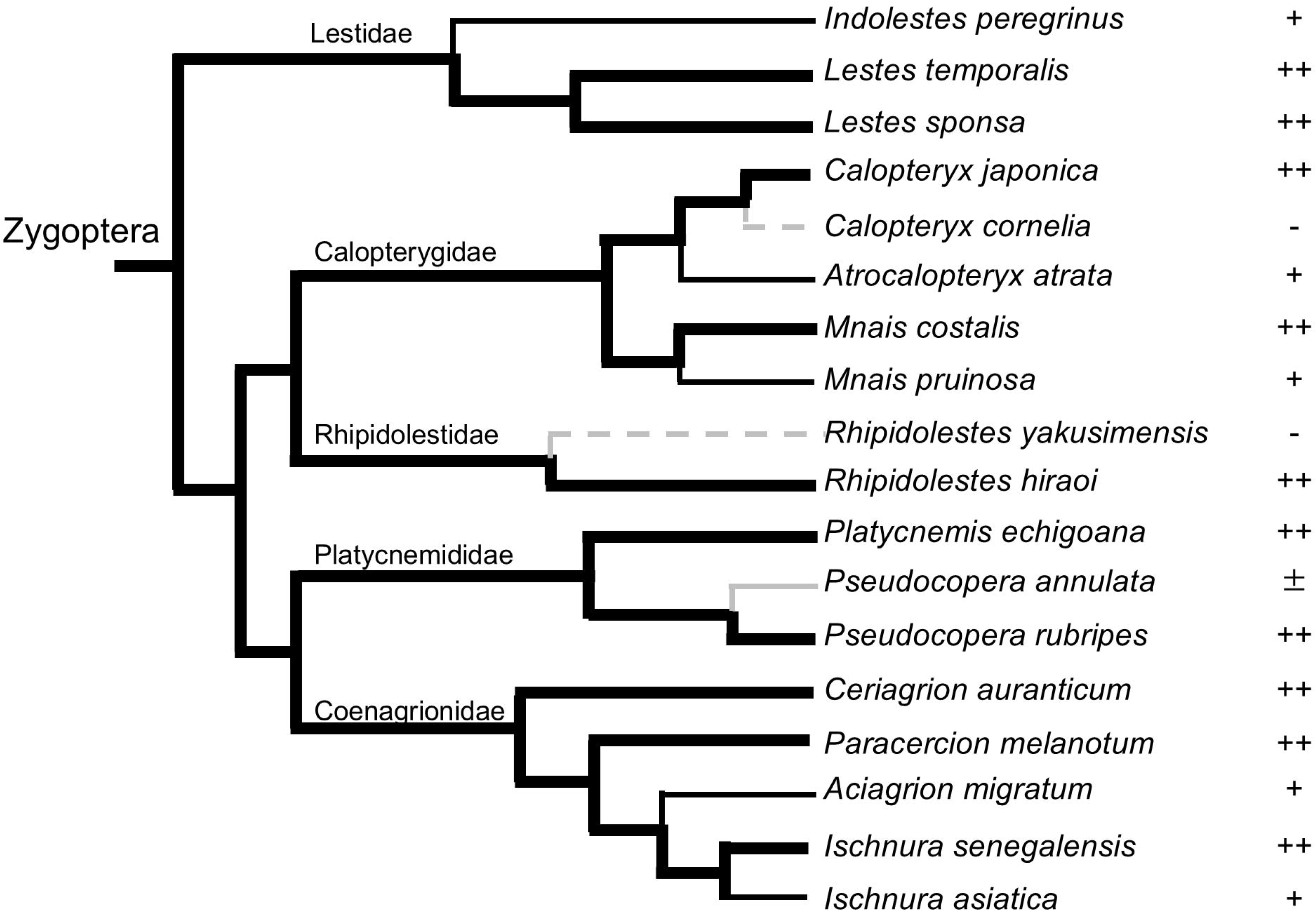
The presence of (TTGGG)_n_ telomeric sequence in damselflies. The odonate phylogeny is based on Ozono et al., 2022. Southern hybridization signal intensity nearly equal to or greater than that of *Orthetrum albistylum* (++, thick black branch), weaker than that of *Orthetrum albistylum* but clearly visible (+, thin black branch); weak signal (+-, gray solid branch); nearly or no detectable signal (gray dotted branch).

The presence/absence of (TTGGG)n was detected within Libellulidae, Corduliidae, Gomphidae, and Aeshnidae in Anisoptera (Figs. 3, 5). and within Calopterygidae, Rhipidolestidae, and Platycnemididae in Zygoptera (Figs. 4, 6), respectively. Moreover, presence/absence of (TTGGG)n was detected within genera *Aeshna* (Aeshnidae), *Sympetrum* (Libellulidae), *Calopteryx* (Calopterygidae), *Rhipidolestes* (Rhipidolestidae), and *Pseudocopera* (Platycnemididae). Of the twelve *Sympetrum* species tested, (TTGGG)n signals were faint excepting *S. bacca* and *S. parvulum*, which is consistent with the TTGGG signal intensity of the chromosome ends by FISH (Figs. 2D, 2E, S3). Overall, these results indicate that the (TTGGG)n is the typical and ancestral telomeric repeat for Odonata, and changes in telomeric repeat sequences have occurred multiple times within families and genera (Figs. 5, 6).

### 3.4. Odonata species possess one or two copies of full-length telomerase reverse transcriptase gene

To investigate whether the lack of (TTGGG)n signals in southern hybridization is related to the loss of the TERT gene, we searched TERT genes in our previously published transcriptome dataset (Futahashi et al., 2015; Futahashi et al., 2019; Okude et al., 2022). We also conducted RNA sequencing for several of the species in which southern hybridization was performed. The full-length TERT gene transcript was detected even in the species which (TTGGG)n southern hybridization signals were absent (*Sympetrum risi*, *Sympetrum pedemontanum*) or faint (*Sympetrum darwinianum, Sympetrum infuscatum, Sympetrum frequens*) (Fig. 7) (accession nos. LC865784–LC865801). We also searched for the TERT gene in dragonflies and damselflies by Blast search to the NCBI genome database, using TERT sequences obtained from our RNAseq data of the same genus or family as queries. The TERT gene was found in all Odonata species surveyed (Fig. 7). Notably, two TERT genes were detected in the transcriptome of Petaluridae species *T. pryeri*, and full-length sequences of two TERT genes, located on the same chromosome in tandem, were detected for other two Petaluridae species, *Tanypteryx hageni* and *Uropetala carovei* (Fig. 7). Phylogenetic analyses suggested that the TERT gene duplication occurred in the common ancestor of *Tanypteryx* and *Uropetala* (Fig. 7). Several partial TERT gene sequences were also detected in the same scaffold as the full-length copy in *Hetaerina titia*, and different scaffolds in *Hetaerina americana*, implying that some Odonata species possesses more than two TERT copies.

**Figure 7.**
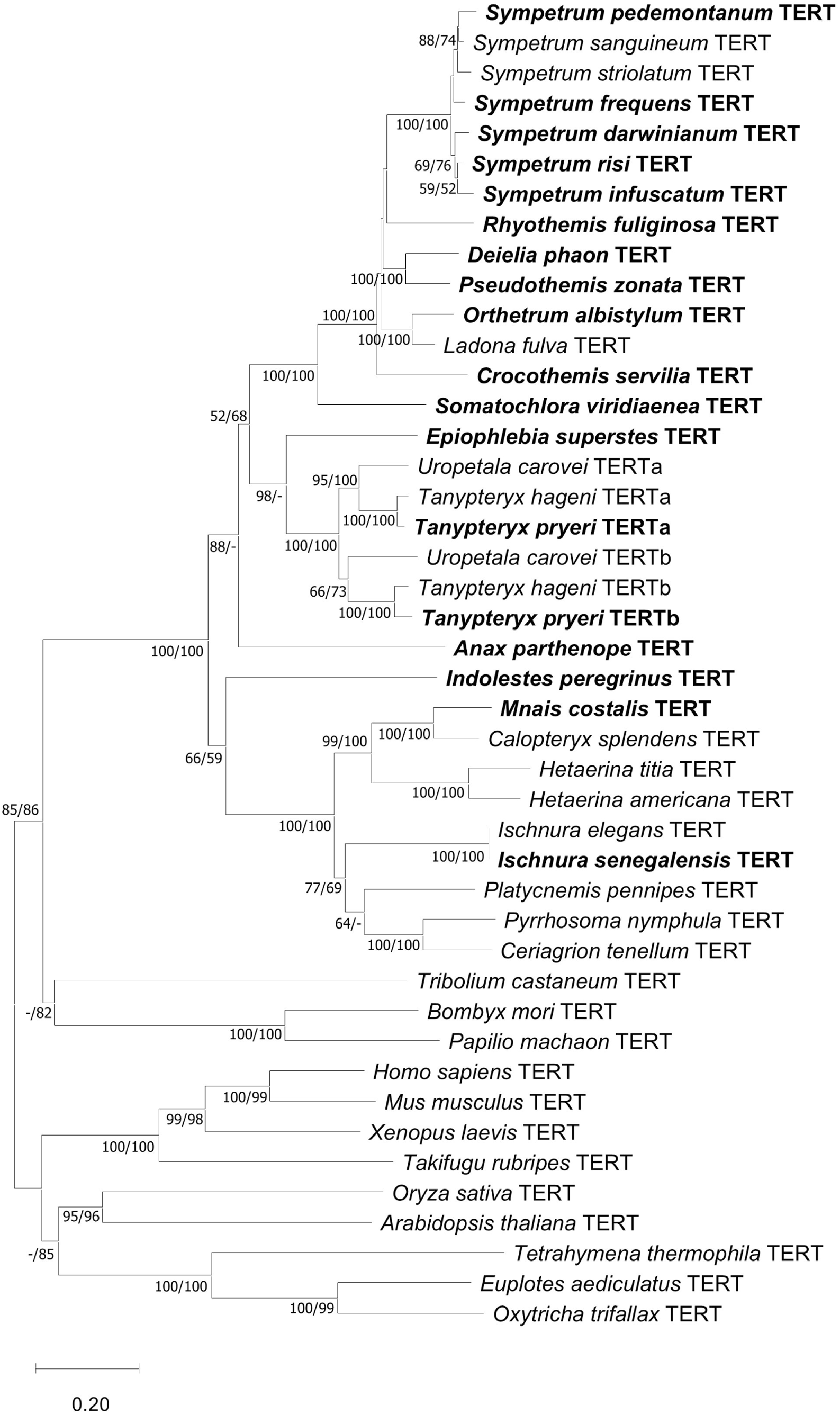
Phylogenetic tree based on the amino acid sequences of the conserved motifs in TERT proteins. The optimal tree constructed by the neighbor-joining method after sequence alignment of the conserved motifs (T, 1, 2, and A through E) of fourteen TERTs. Odonata TERT sequence identified by RNAseq are indicated in bold. Statistical support values for each clade are indicated in the order of bootstrap probability of the neighbor-joining analysis and bootstrap probability of the maximum-likelihood analysis from left to right, in which hyphens indicate values less than 50%.

## 4. Discussion

In this study, we found that (TTGGG)_n_ is a typical and ancestral telomeric repeat sequence in the order Odonata by combination of genomic analysis, FISH, and Southern hybridization. We confirmed that the majority of Odonata species covering all three suborders have (TTGGG)n as the telomere repeat sequence.

In FISH analyses, we recognized chromosomes looped in the same direction in damselflies *Lestes temporalis* and *Ischnura asiatica*, and dragonflies *Anotogaster sieboldii* and *Crocothemis servilia* (Figs. 2H, S2). Their chromosome configuration strongly resembled the telomere bouquet arrangement. In vertebrates and yeasts, the telomere bouquet is observed during meiotic prophase (Mytlis et al., 2023), in which the telomeres form a cluster beneath the nuclear envelope. Since chromosome configuration resembling telomere bouquet arrangement has been also observed from Giemsa stains of testes of dragonflies *Somatochlora metallica* (Nokkala et al., 2002) and *Orthemis ambinigra* (Mola et al., 2021), formation of telomere bouquet structure is a common characteristic of male meiotic prophase in Odonata. This is in contrast with the Rabl configuration of *Drosophila* meiotic prophase during which centromeres instead of telomeres are tethered to the nuclear envelope (Mytlis et al., 2023), and the *Bombyx* male meiosis prophase during which only one end of the chromosomes is tethered to the nuclear envelope (Rosin et al., 2021), indicating that chromosome configuration in meiosis differs among insects.

It should be noted that the species with weak or no (TTGGG)n signals were found in several lineages within the Anisoptera and Zygoptera. This indicates that the alteration of telomere sequences has repeatedly occurred in Odonata. Repeated changes in telomeric repeat sequences have been reported in Coleoptera and Hymenoptera (Prušáková et al., 2021; Zhou et al., 2022; Lukhtanov and Pazhenkova, 2023). While the diversification of telomeric repeat sequences of Coleoptera has been hypothesized to be correlated with its species-richness and diversity (Prušáková et al., 2021), it is noteworthy that repeated changes in telomere repeat sequences have occurred in Odonata, despite only 5,500 species are extant in this order.

In the genus *Sympetrum*, the signal intensity for (TTGGG)n varied between species. Of the twelve *Sympetrum* species tested for Southern hybridization, only *S. bacca* and *S. parvulum* gave relatively strong (TTGGG)n signals, which is consistent with FISH signal intensity of *S. frequens*, *S. bacca* and *R. fuliginosa* (Figs. 2D, 2E, S3). Considering the phylogenetic relationships of *Sympetrum* (Pilgrim and von Dohlen, 2012; Ozono et al., 2022), the results of Southern hybridization suggest that alterations in the telomeric (TTGGG)n repeats have repeatedly occurred even within the genus *Sympetrum* (Fig. 5). In this context, it is interesting to note the higher sequence divergence (6 to 13%) of the (TTGGG)n sequence in *S. striolatum* (Supplementary data 3), although it cannot be confirmed whether these sequences are located at chromosome ends or not. As we could not identify candidates for telomeric repeat sequences other than (TTGGG)n, identification of the alternative telomeric repeats in Odonata awaits the telomere-to-telomere genome assembly in various species.

In the Petaluridae species *Tanypteryx pryeri*, only a faint signal was observed in genomic Southern hybridization using the (TTGGG)n probe, which suggested the alteration of telomeric sequence in the species. In the NCBI deposited genome sequence of the closely related species *Tanypteryx hageni*, (TTGGG)n was detected in several chromosomes but not at the termini of the chromosome. Notably, two copies of the TERT gene were present in Petaluridae species (Fig. 7). Since telomeric repeats are dependent on the sequence of the telomerase RNA subunit, the presence of multiple TERT genes may be an accelerator for diversification and evolution of telomeric repeats, which deserves further studies.

## Acknowledgements

We thank Shigeru Matsubara, Mitsutoshi Sugimura, Akira Ozono, Taku Kitayama, Bin Hirota, Akiyoshi Nakata, Yota Mizuno, Genta Okude, Takema Fukatsu, Kazuo Yoshida, Mitsuhiro Fuwa, Yoshiaki Sakai, Tatsuya Nakata, Kazunari Hisada, Michihiko Takahashi, Tomohito Noda, Yuta Higuchi, Hiroyuki Futahashi, Masafumi Futahashi, and Naoki Futahashi for Odonata specimens. Silkworm strain p50 was provided by the National BioResource Project (NBRP) of Japan. We thank Yusuke Kobayashi for the use of CCD camera LAS-3000.

## CRediT authorship contribution statement

Tatsuhiro Gotoh: Writing – original draft, Writing – reviewing and editing, Visualization, Validation, Resources, Methodology, Investigation, Formal analysis

Haruka Suzuki: Writing – original draft, Visualization, Validation, Methodology, Investigation

Minoru Moriyama: Writing – reviewing and editing, Formal analysis, Data curation

Ryo Futahashi: Writing – original draft, Writing – reviewing and editing, Visualization, Resources, Investigation, Formal analysis, Funding acquisition, Data curation

Mizuko Osanai-Futahashi: Writing – original draft, Writing – reviewing and editing, Visualization, Supervision, Resources, Project administration, Investigation, Formal analysis, Funding acquisition, Conceptualization

## Data availability

RNA-seq data was submitted to DDBJ under accession numbers DRR016606-DRR016611, DRR016624-DRR016629, DRR022340-DRR022342, DRR082407-DRR082410, DRR656569-DRR656589. The nucleotide sequences of TERT genes were submitted to DDBJ under accession numbers LC865784–LC865801.

