## Supplementary Figures for "Acquisition and repeated alteration of (TTGGG)_n_ telomeric repeats in Odonata (dragonflies and damselflies)"

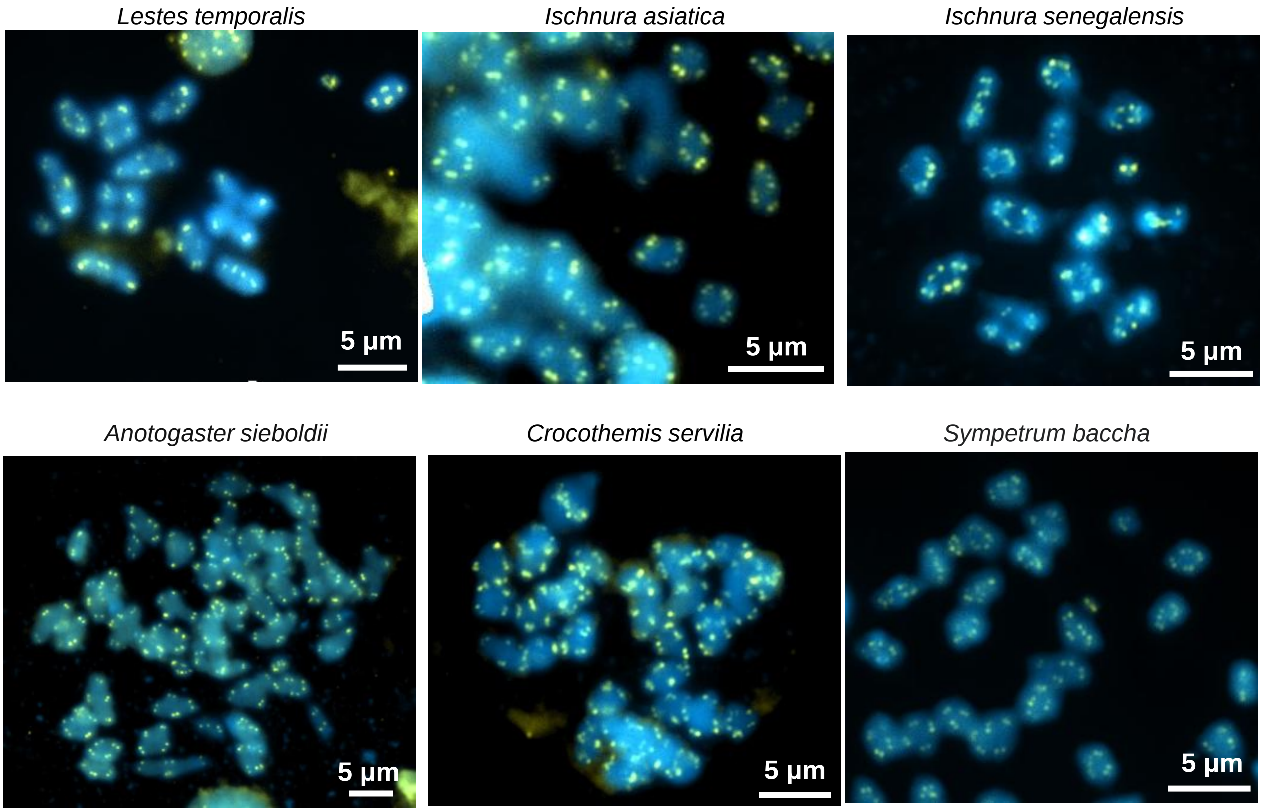


**Fig. S1 TTGGG signals detected in diakinesis of Odonata species by Fluorescence *in situ* hybridization.**

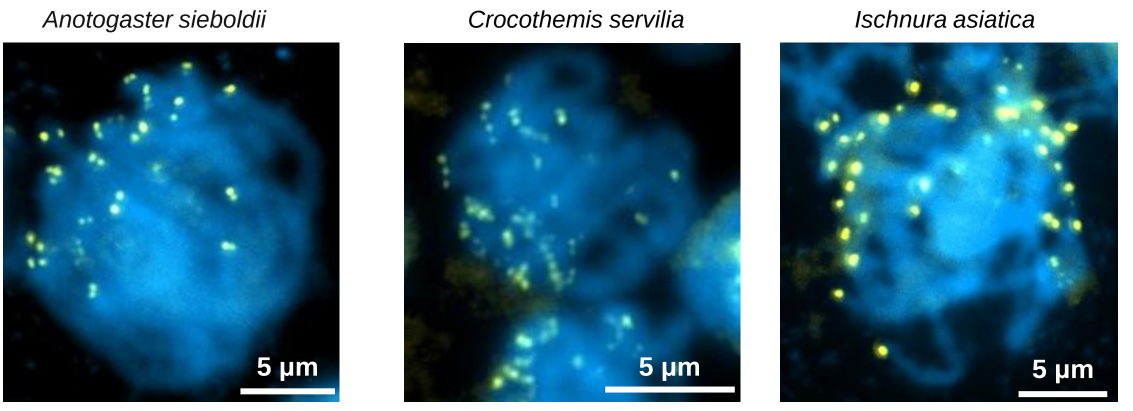


**Fig. S2 Telomere bouquet observed in Odonata species.**
Signals of TTGGG probes are indicated in yellow, and chromosomes were counterstained with DAPI or PI.

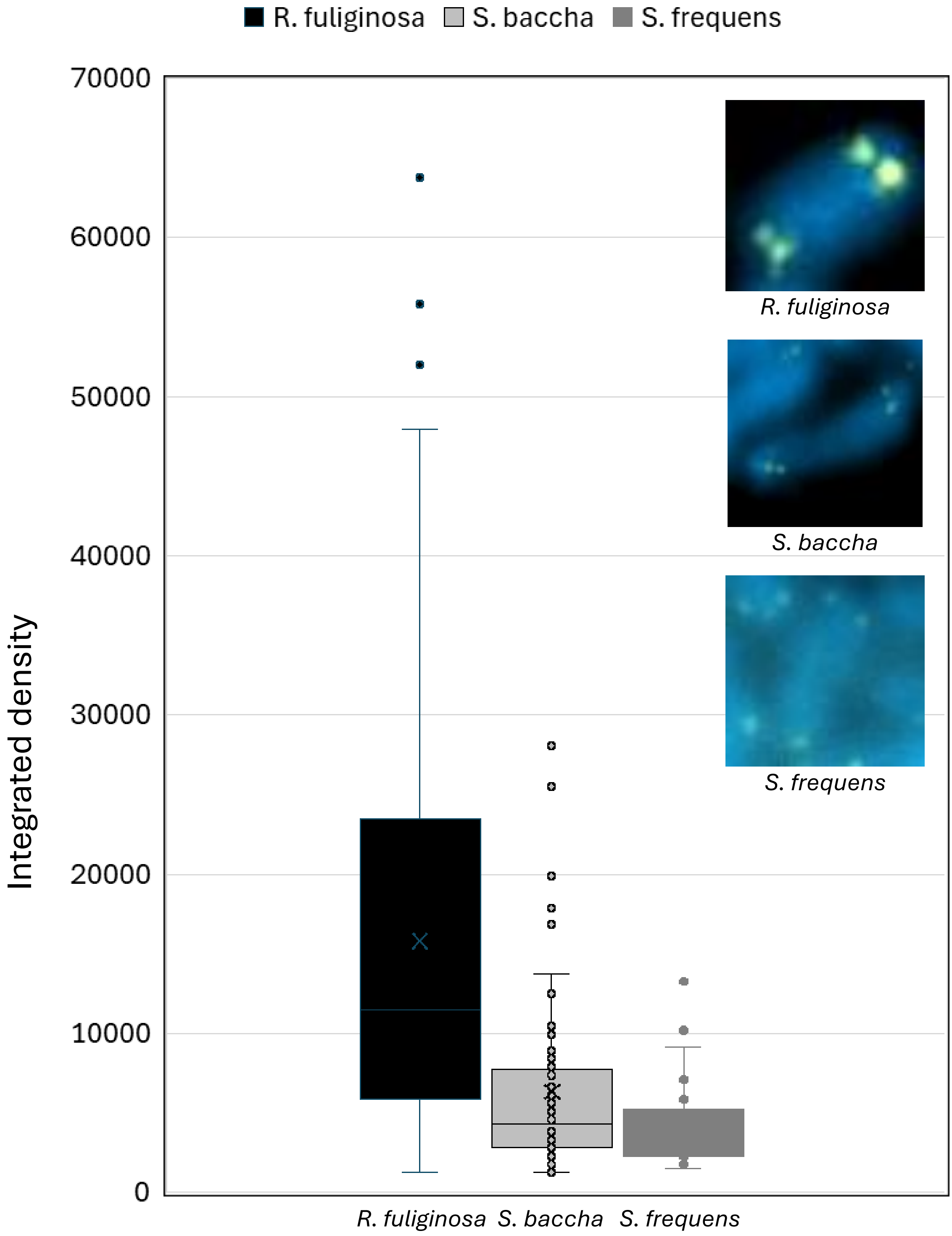


**Fig. S3 Comparison of signal intensity of (CCCAA)3CCC PNA probe.**
Comparison of signal intensity of (CCCAA)3CCC PNA probe between *S. frequens*, *S. baccha*, *R. fuliginosa*. Images taken under the same exposure conditions, and the integrated density of fluorescent signals (>5) for (CCCAA)3CCC PNA probe were analyzed by ImageJ.

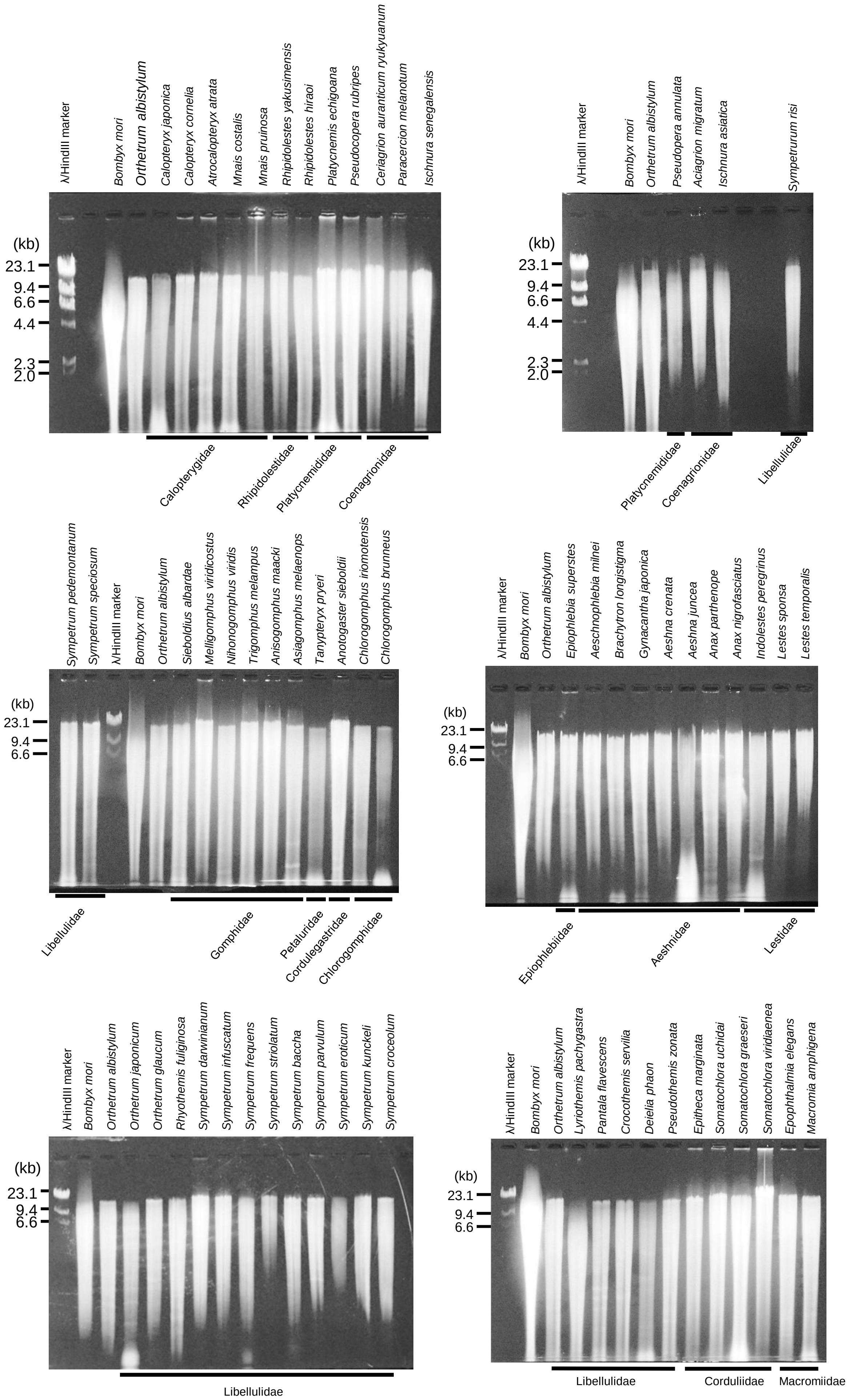


**Figure S4. Electrophoresis pattern of Hind III digested genomic DNA for southern hybridization prior to membrane transfer.**

Appoximately 1 micrograms of HindIII digested genomic DNA were applied for each lane.
